# HSF1 excludes CD8+ T cells from breast tumors via suppression of CCL5

**DOI:** 10.1101/2022.05.12.491688

**Authors:** Curteisha Jacobs, Sakhi Shah, Wen-Cheng Lu, Haimanti Ray, John Wang, George Sandusky, Kenneth P. Nephew, Xin Lu, Sha Cao, Richard L. Carpenter

## Abstract

Heat shock factor 1 (HSF1) is a stress-responsive transcription factor that promotes cancer cell malignancy. A novel HSF1 Activity Signature (HAS) was found to be negatively associated with antitumor immune cells in breast tumors. Knockdown of HSF1 decreased tumor size and caused an influx of several antitumor immune cells, most notably CD8+ T cells. Depletion of CD8+ T cells prevented tumors from shrinking after knockdown of HSF1, suggesting HSF1 prevents CD8+ T cell influx to avoid immune-mediated tumor killing. HSF1 was also found to suppress expression of CCL5, a chemokine for CD8+ T cells, that significantly contributed to the attraction of CD8+ T cells upon the loss of HSF1. This study demonstrates a model whereby HSF1 suppresses CCL5 leading to reduced CD8+ T cells in breast tumors that prevented immune-mediated destruction. For the first time, these studies indicate HSF1 suppresses antitumor immune activity within tumors.

## Introduction

Breast cancer is the second leading cause of cancer related deaths in women with 1 in 8 developing invasive breast cancer over the course of their lifetime (1). Immune checkpoint therapy in breast cancer patients has had mixed results with triple-negative breast cancer (TNBC) patients primarily being the beneficiary of this therapeutic approach (2,3). Breast cancer has historically thought to be a low immunogenic tumor (4,5). TNBC has largely benefitted from immune checkpoint therapy due to it being the small subset of breast cancer that has shown to have PD-L1 expression and higher lymphocytic infiltration relative to other breast cancer subtypes (6–9). While PD-L1-targeting checkpoint therapy was approved for high risk TNBC, only a small percentage of these patients respond to immune checkpoint inhibition prompting the approval for anti-PD-L1 therapy to be removed for this patient population in 2021. Previous studies have identified that higher lymphocyte infiltration in breast cancers is associated with improved survival and improved response to treatments (10–13). Infiltration of CD8+ T cells, the primary cytotoxic lymphocyte, is an independent predictor of patient outcome and these immune cells are trafficked to tumors through chemotactic cytokines (14). While low tumor mutational burden (TMB) is a significant factor in the low lymphocyte infiltration in breast cancer, TMB does not fully explain low lymphocyte infiltration, indicating that additional mechanisms suppress lymphocyte attraction to breast tumors.

Heat Shock Factor 1 (HSF1) is a master transcription factor regulator of the heat shock response. The classical function of HSF1 is to regulate expression of chaperone genes in response to cellular stressors (15). HSF1 can be hyperactivated due to increased proteotoxic stress and upregulation of activating kinases including AKT1, mTORC1, and p38 among others (15–18). However, the seminal finding of a unique transcriptional program of HSF1 in cancer cells that is distinct from the heat stress response transcriptional program indicated that HSF1 has additional functions in cancer (19). It is now recognized that HSF1 plays key roles in cancer cells by promoting several pro-tumor processes including epithelial-to-mesenchymal transition, promotion of the cancer stem-like state, promotion of cancer-associated metabolic changes, and several others that all support the malignancy of cancer cells (15–17,19,20). Furthermore, HSF1 levels and activity have been associated with worsened patient outcomes in many cancer types, including breast cancer (17,19,21). However, the role of HSF1 in tumor-immune interactions and avoiding immune destruction remains unclear.

Gene signatures, or sets of genes that represent biological functions or features, have gained popularity since the introduction of gene set enrichment analysis (GSEA) and the molecular signatures database (mSigDB) (22). Here we report a new gene signature termed “HSF1 Activity Signature,” or HAS, a 23-gene signature that reports HSF1 transcriptional activity generated using a novel approach. By using the HAS, we uncovered a novel mechanism underlying breast tumor evasion of immune destruction based on inhibition of CD8+ T cell recruitment to tumors by HSF1-mediated suppression of CCL5, a key chemokine regulating CD8+ T cells.

## Results

### Identification of an HSF1 Activity Signature (HAS) that detects changes in HSF1 transcriptional activity

HSF1 was originally discovered for its role as the master regulator of the heat shock response (15,17). The discovery of new functions of HSF1 has particularly been fruitful in the context of cancer where it has been found to regulate a target gene set distinct from the genes targeted by HSF1 in response to heat shock and, consequently, play a role in many processes that promote malignancy (15,17,19). In this study, we utilized a new computational approach to associate HSF1 transcriptional activity with cancer-related processes and patient outcomes. HSF1 has a complex post-translational activation process whereby it must localize to the nucleus in order to undergo trimerization and DNA binding, in addition to phosphorylation prior to recruiting general transcription factors to initiate transcription at target genes. Because of this complex activation process at the protein level, RNA levels of the HSF1 gene are not highly predictive for HSF1 transcriptional activity and HSF1 RNA does not significantly increase expression in response to heat shock (Suppl Fig. 1A-C). We reasoned that the most predictive gene set for HSF1 transcriptional activity will be RNA expression levels of genes that 1) are direct HSF1 target genes, 2) have high intra-gene correlation within the gene set across multiple datasets, 3) decrease expression when HSF1 is knocked down or inhibited, 4) increase expression in response to heat shock, and 5) show increased expression in cancer samples compared to normal. Therefore, we attempted to identify a set of genes that meet these criteria utilizing the gene selection procedures outlined in Figure 1A.

**Figure 1:**
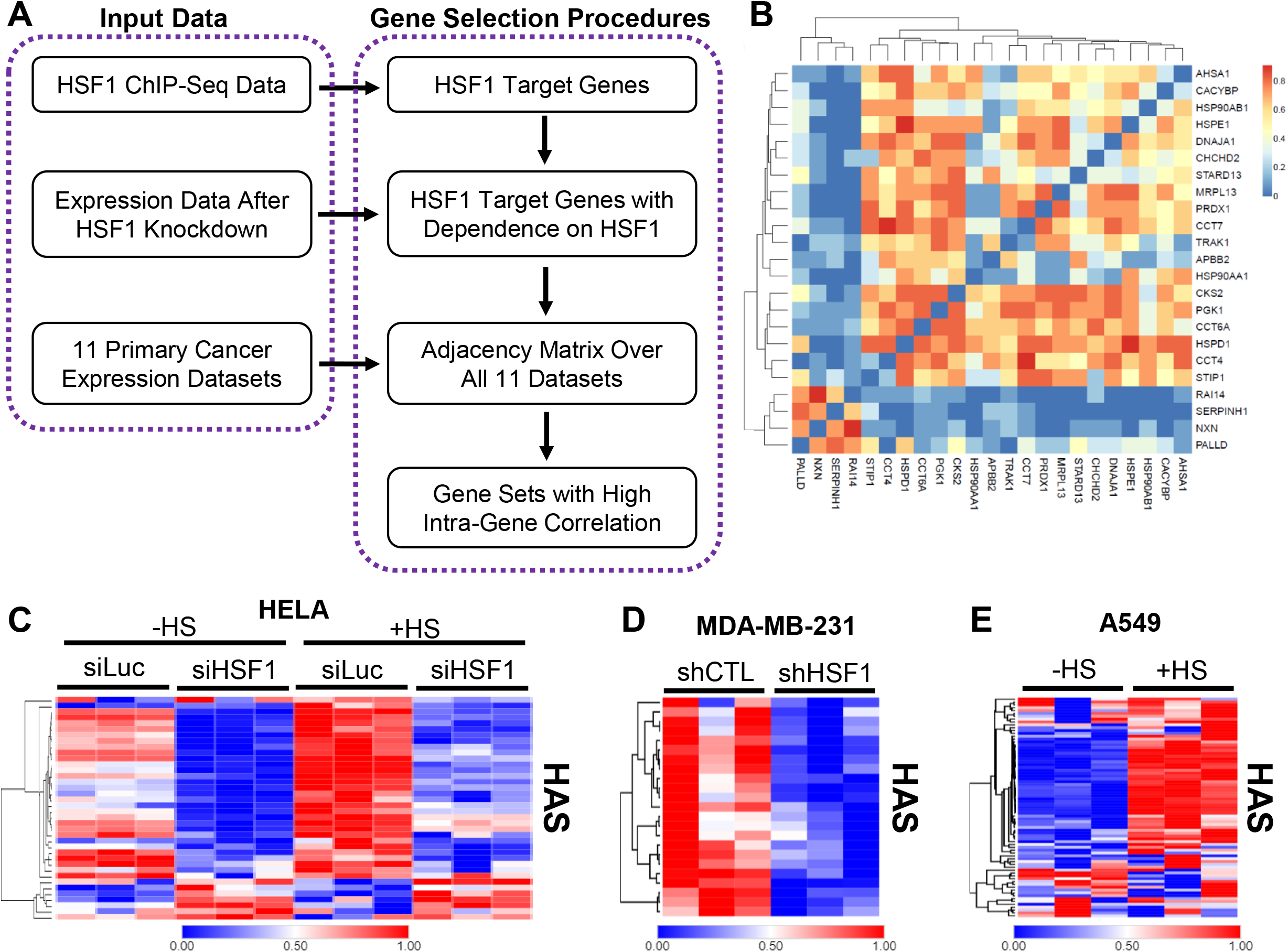
Identification of the HSF1 Activity Signature (HAS). A) The schema for identifying an HSF1 activity signature, which included input data from ChIP-Seq data to identify direct targets followed by removing genes not dependent on HSF1 for their expression and finally generating gene sets that have high intragene correlation. B) Correlation matrix of the 23-gene HAS across 11 cancer datasets. C-E) Heat maps were generated from publicly available expression data from Hela cells with HSF1 knockdown and/or heat stress (C), MDA-MB-231 cells with or without HSF1 knockdown (D), and A549 cells with or without heat stress (E).

We first identified genes that are direct transcriptional targets of HSF1 utilizing more than 40 ChIP-Seq samples in the public domain (19,23,24). All unique genes were included in the initial gene list as potential HSF1 target genes. We then removed genes whose expression was not decreased with the knockdown of HSF1 based on differential expression analysis. We then conducted an integrated co-expression analysis for the remaining genes using 11 different cancer expression datasets (Suppl. Table 2). This co-expression analysis resulted in identification of 9 unique gene sets for consideration (Suppl. Table 4). One gene set, hereafter referred to as the HSF1 Activity Signature (HAS), was found to have the highest intra-gene correlation (Fig. 1B) and outperformed all other gene sets in detecting changes in HSF1 activity after HSF1 knockdown or response to heat stress (Fig. 1C, Suppl. Fig. 1D-K). The HAS gene set was consistently decreased when HSF1 was knocked down or when HSF1 was inhibited with DTHIB (Fig. 1C-D; Suppl. Fig.2A-C). Inversely, the HAS increased after heat stress (Fig. 1C-E; Suppl. Fig. 2D-E). Principal component analysis (PCA) showed that PC1 of the HAS accounted for 60-85% of the variance across these experiments (Suppl. Fig. 3A-G). To statistically compare HAS across these groups, PC1 scores for each sample was compared and showed a significant decrease in HAS with HSF1 knockdown while a significant increase was observed in samples with heat shock (Suppl. Fig. 3A-G). Genes associated with the HAS were analyzed with gene ontology that revealed several ontologies involved with protein folding, protein stabilization, heat shock proteins, unfolded proteins, chaperones, chaperonins, and chaperone complexes that were enriched (Suppl Fig. 4A-C) and indicative of the known function of HSF1 in proteostasis. Heat shock elements (HSEs), the HSF1 binding motif, were also the most enriched motif among the genes within the HAS (Suppl. Fig 4D). These data indicate the 23 gene set HAS can reliably detect changes in HSF1 transcriptional activity.

### HSF1 activity is associated with breast cancer patient outcomes and molecular characteristics

HSF1 has previously been associated with several cancer phenotypes and increased expression and transcriptional activity in cancer cells (15,17). The HAS genes showed a clear increase in expression in tumor samples compared to matched normal adjacent tissue (Fig. 2A), indicating the HAS was able to detect an increase in HSF1 activity in tumors. We next assessed whether the HAS could serve as a biomarker for outcomes of breast cancer patients as previous studies show active HSF1 was associated with worse outcomes (19,21,23). As shown in Figs. 2B-C, high HAS was associated with worse overall survival (OS) in the TCGA breast cancer cohort and the METABRIC breast cancer cohort whereas expression of the HSF1 gene alone was not as consistent in reproducing this association (Suppl. Fig. 5A-B). In addition to Kaplan Meier and Log Rank tests, age-adjusted hazard ratio for HAS was also significantly associated with worse OS in the TCGA (HR=2.42, 95% CI: 1.62-3.62; p<0.001) and the METABRIC (HR=1.58, 95% CI: 1.33-1.88; p<0.001) cohorts. HAS was also associated with worse metastasis-free survival of breast cancer patients (Fig. 2D) whereas expression of the HSF1 gene was not significantly associated with metastasis-free survival (Suppl. Fig. 5C), further supporting previous studies suggesting a potential role for HSF1 in metastasis (19,20,25). Taken together, these results indicate that HAS can function as a marker of HSF1 activity and HAS predicted breast cancer patient outcomes consistent with previous assessments for HSF1 activity that include measurement of activity by nuclear HSF1 levels or S326 phosphorylated HSF1 (19–21).

**Figure 2:**
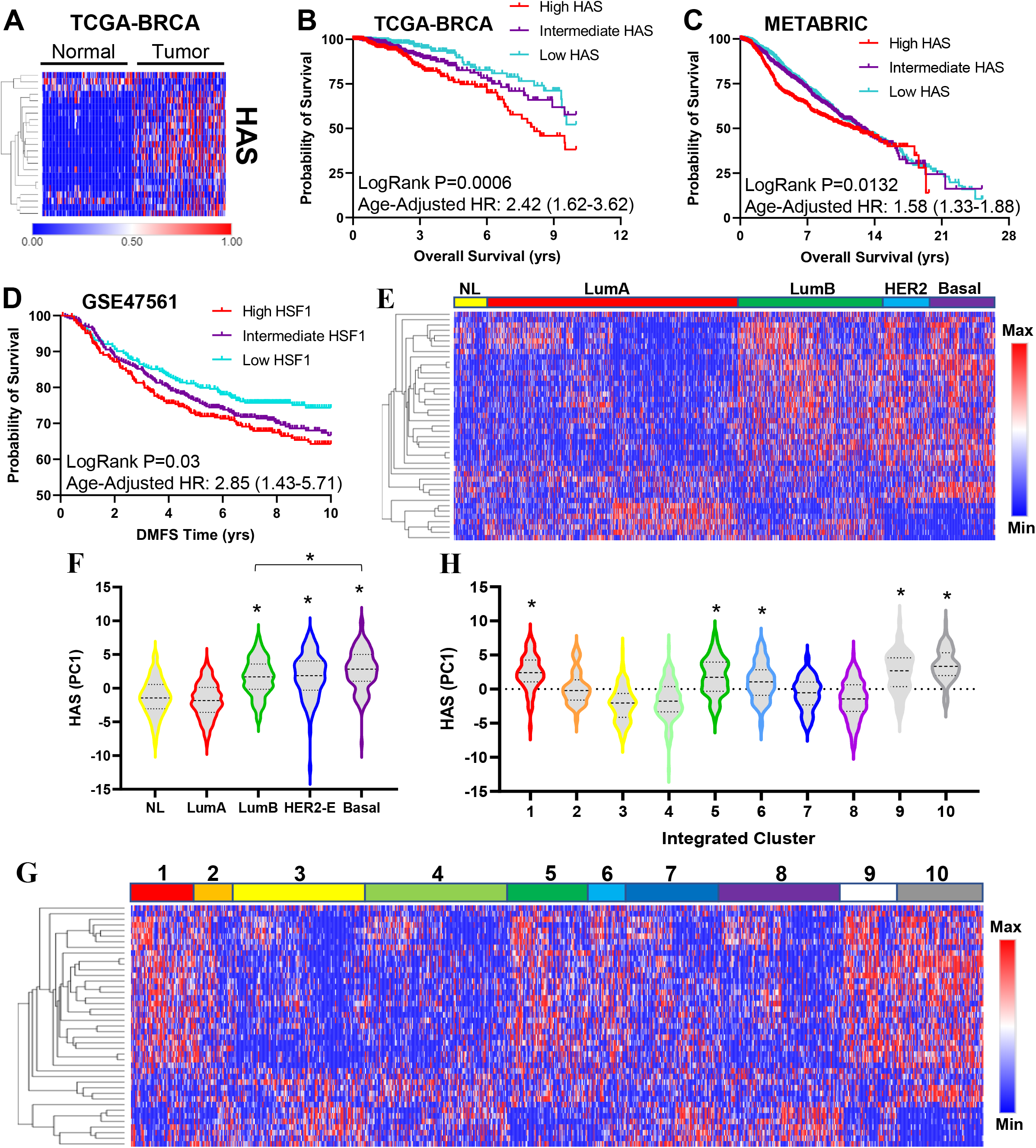
HAS is associated with breast cancer outcomes and molecular subtypes. A) Heat map was generated using matched adjacent normal and tumor expression data from the TCGA-BRCA cohort (n=100). B-D) Patients in the TCGA-BRCA (B), METABRIC (C), and GSE47561 (D) cohorts were sorted by their HAS scores and Kaplan-Meier plots were generated for overall survival (B-C) or metastasis-free survival (D). E-F) Heat map (E) was generated for the HAS of the METABRIC cohort delineated by subtype and HAS PC1 scores (F) for each subtype were compared across subtypes via one-way ANOVA with Tukey’s post-hoc test. *Indicates significance compared to Normal-Like. G-H) Heat map (G) was generated for the HAS of the METABRIC cohort delineated by METABRIC Clusters and PC1 scores (H) were plotted across METABRIC Clusters. *Indicates significance compared to the lowest cluster (#3). NL=Normal-Like; LumA=Luminal A; LumB=Luminal B; HER2-E=HER2-enriched; HAS=HSF1 activity signature.

We next assessed the relationship of the HAS with molecular subtypes of breast cancer. HAS was increased within the Luminal B (LumB), HER2-enriched (HER2-E), and Basal subtypes in both the METABRIC and TCGA cohorts compared to the Normal-Like (NL) and Luminal A (LumA) subtypes (Fig. 2E-F, Suppl. Fig. 5D-E). The HAS was also assessed in the METABRIC Integrated Clusters (IntClust) where the HAS was upregulated in several integrated clusters that appear to mirror HAS activation from molecular subtypes. HAS was increased in IntClust 10, which is mostly associated with the basal molecular subtype, and IntClust 5, which is closely associated with the HER2-enriched molecular subtype (Fig. 2G-H). Interestingly, HAS was also elevated in clusters 1, 6, and 9 that are primarily ER-positive cancers that includes overlap with the LumB molecular subtype. Upregulation of the HAS in these clusters likely points towards a recently-identified interaction of HSF1 with ERα in breast cancer (26) and an interaction between HSF1 and HER2 signaling, which has been established (16,25,27). Furthermore, HAS was associated with both disease-free survival and overall survival across the spectrum of TCGA cancer types (Fig. 3A-B). HSF1 has previously been associated with outcomes for many cancer types and the HAS reflected many of these known associations including liver (28), lung (29), melanoma (23), esophageal (30), and head/neck (31) cancers among others.

**Figure 3:**
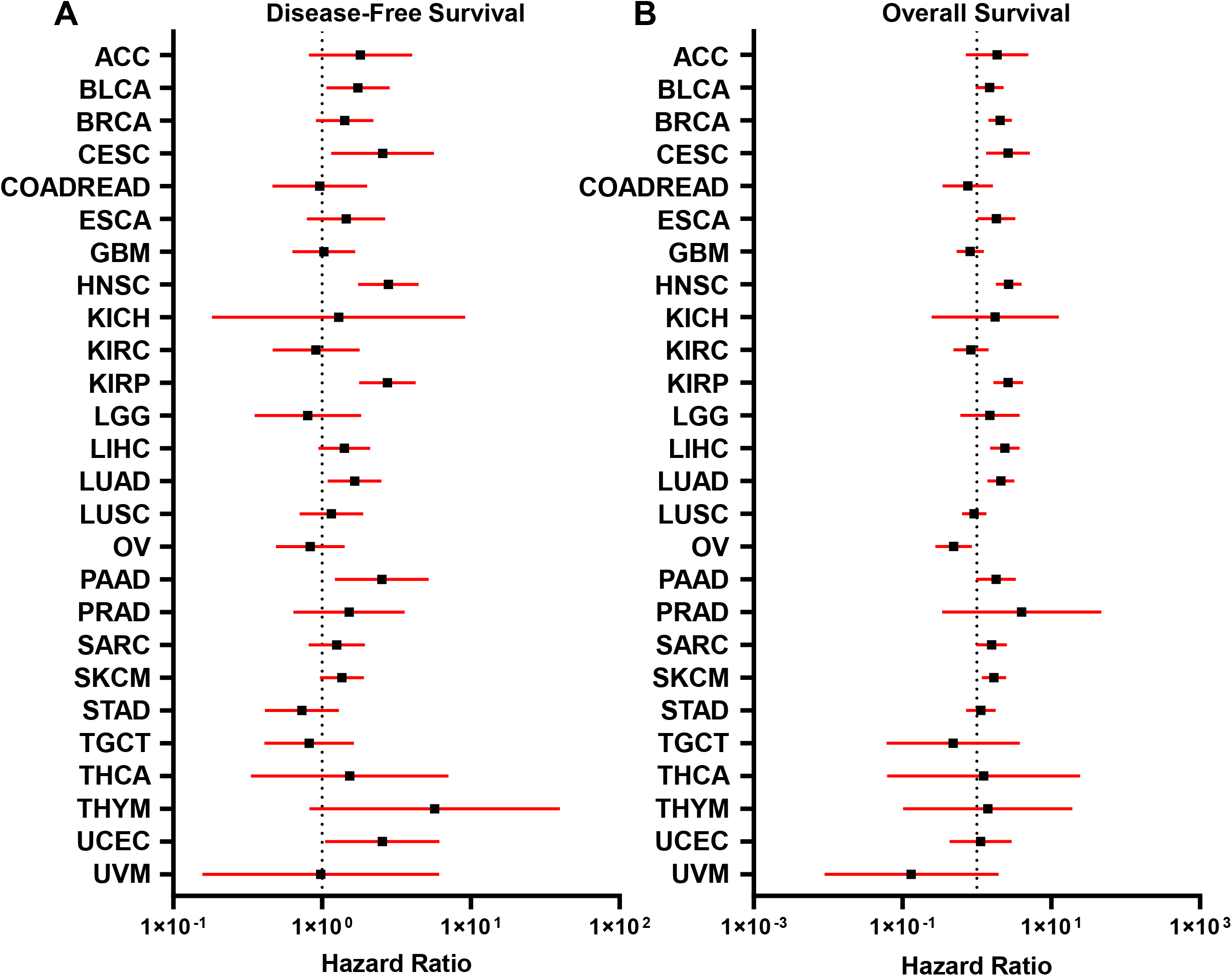
Association of HAS with Outcomes Across TCGA Cancer Types. A-B) Cox hazard ratios were calculated for HAS, adjusted for age, across the TCGA cancer types for Disease-Free Survival (A) and Overall Survival (B). Forest plots were generated with hazard ratios (black square) with 95% confidence intervals (red bars).

### HSF1 Activity has a Negative Association with the Presence of CD8+ T cells in Breast Tumors

Previous reports indicated a possible association between HSF1 and tumor-immune interactions as well as immune function (19,32). To investigate the relationship between HSF1 activity and tumor-immune interactions, the HAS was used to sort tumors from breast cancer patient cohorts in the TCGA-BRCA (n=1100), METABRIC (n=1996), and GSE47561 (n=1570) into high and low HSF1 activity groups. These groups were then subjected to gene set enrichment analysis (GSEA) to determine association of HSF1 activity with immune cell populations using multiple immune cell estimation algorithms (33–38). Interestingly, HSF1 activity was negatively associated with the presence of several immune cell types including CD8+ T cells, CD4+ T cells, and B cells across several different signatures for these cell types (Fig. 4A). Utilizing the immune cell estimation algorithms, it was consistently observed that patients with high HAS scores showed lower CD8+ T cells proportions compared to patients with low HAS scores (Fig. 4B-C, Suppl. Fig. 6A-B). To determine if this relationship between HSF1 and CD8+ T cell proportions is unique to any breast cancer subtype, GSEA was performed on subpopulations of patients with different receptor status. The relationship between the HAS score and CD8+ T cells appeared to be stronger in triple-negative breast cancer (TNBC) compared to tumors with other receptor statuses (Fig. 4D-F, Suppl. Fig. 6C-G). Breast cancer patients who had high HAS scores and low CD8+ T cell proportions showed significantly worse overall survival or metastasis-free survival compared to patients with lower HAS and higher CD8+ T cell proportions, which was consistent in the TNBC population (Fig. 4G-H, Suppl Fig. 6H-K).

**Figure 4:**
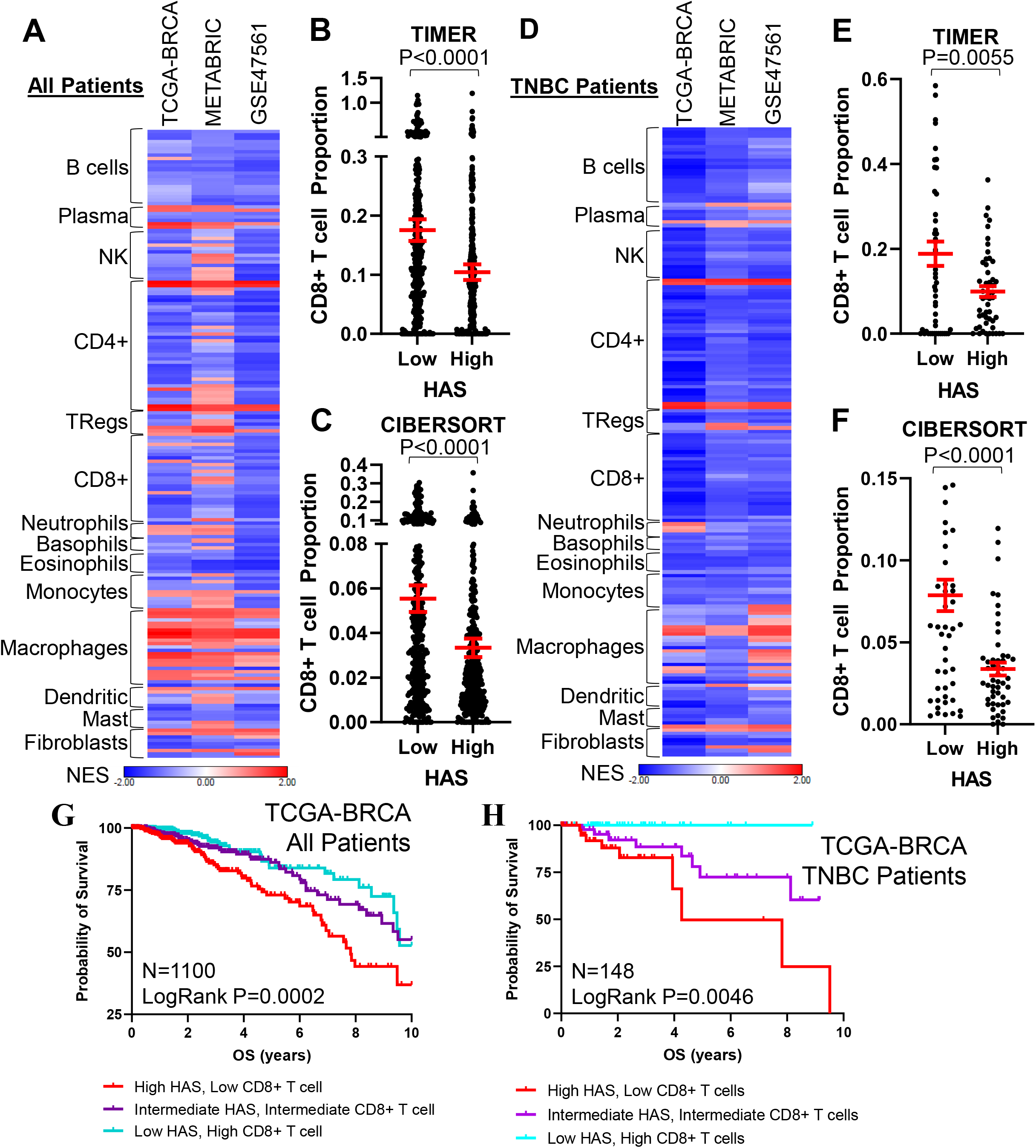
HAS is Negatively Associated with Presence of CD8+ T cells. A) GSEA was performed in TCGA-BRCA, METABRIC, and GSE47561 cohorts with patients separated into high or low HAS scores. Signatures for immune cell types were assessed for enrichment with high or low HAS patients. Normalized enrichment scores (NES) are plotted on a heat map. B-C) CD8+ T cell proportions were estimated in the TCGA-BRCA cohort using TIMER (B) and CIBERSORT (C). CD8+ proportions are plotted in high and low HAS patients. D) GSEA was performed as in (A) in only TNBC patients in the TCGA-BRCA cohort. E-F) CD8+ T cell proportions were estimated in TNBC patients in the TCGA-BRCA cohort using TIMER (E) and CIBERSORT (F). CD8+ proportions are plotted in high and low HAS patients. G-H) Patients in the TCGA-BRCA cohort were separated by HAS scores and CD8+ T cell proportions estimated by CIBERSORT and Kaplan-Meier graphs were plotted for patient outcomes using all patients (G) or only TNBC patients (H). NK=Natural killer cells; Tregs=T regulatory cells; HAS=HSF1 activity signature.

These data suggest that tumors with high HSF1 activity have lower CD8+ T cells. To confirm these observations, tumor tissues from a cohort of 114 breast cancer patients spanning all subtypes were subjected to IHC for the active mark of HSF1 (phosho-S326) that allowed us to separate tumors into high or low HSF1 active tumors based on nuclear presence of p-HSF1 (Fig. 5A-B). These same specimens were co-stained with CD3A and CD8A antibodies to identify the CD8+ T cells. Similar to the computational analyses, these patient specimens had significantly less CD8+ T cells in tumors with high levels of active HSF1 (Fig. 5C-D). This relationship was maintained in TNBC patient tumors but was not significant in ER+ or HER2+ patient tumors (Fig. 5E-F, Suppl. Fig. 7A-D). These data indicate a consistent relationship wherein tumors with high HSF1 activity have lower levels of CD8+ T cells.

**Figure 5:**
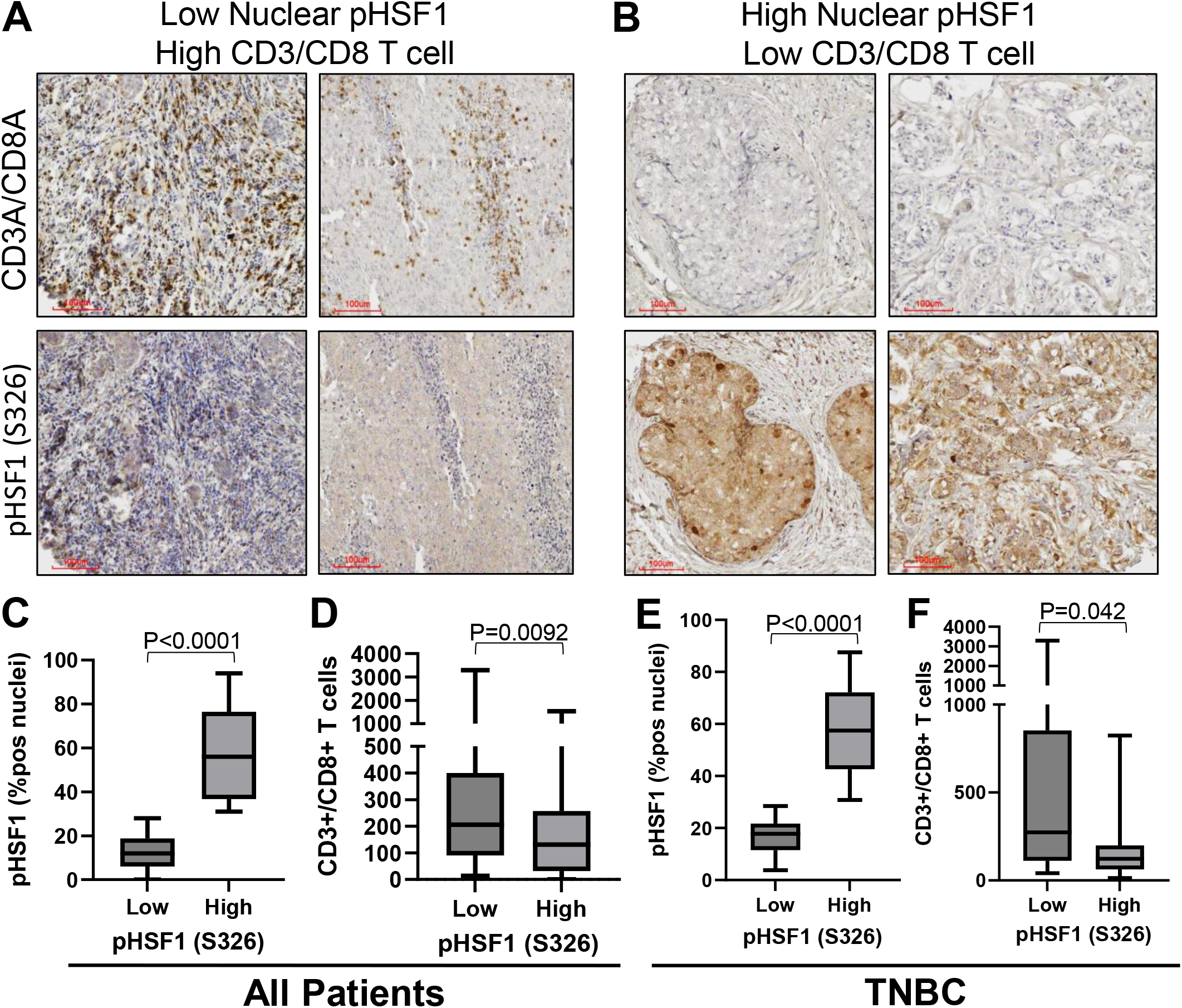
Active HSF1 in Breast Cancer Tumor Specimens Coincides with Low CD8+ T cells. A-B) A cohort of 114 breast tumors were subjected to IHC with antibodies for CD3A, CD8A, and pHSF1 (S326). C-D) All patients (n=114) were separated into high (n=46) or low (n=68) HSF1 active tumors based on nuclear positivity percentage for pHSF1 (C) and CD8+ T cells (D) were compared between these two groups based on active HSF1 levels using a student’s t-test. E-F) Only triple-negative breast cancer (TNBC) patients (n=38) were separated into high (n=21) or low (n=17) HSF1 active tumors based on nuclear positivity percentage for pHSF1 (E) and CD8+ T cells (F) were compared between these two groups based on active HSF1 levels using a student’s t-test.

### HSF1 Functionally Affects the Level of CD8+ T cells in Breast Tumors

Having demonstrated a significant negative association between HSF1 activity and the presence of CD8+ T cells using computational approaches and assessment of patient specimens, it was important to examine where a functional relationship also exists. First, to test for a functional effect of HSF1 on CD8+ T cells, 4T1 cells were engineered to express control or HSF1-directed shRNA (Suppl. Fig. 8A). These cells were orthotopically grown in the mammary glands of Balb/c mice for 3 weeks, after which it was confirmed that HSF1 expression was lost and resulted in a significantly smaller tumor volume (Fig. 6A, Suppl. Fig 8B). The number of CD8+ T cells in these tumors were assessed with IHC that showed HSF1 knockdown tumors had significantly more CD8+ T cells compared to control tumors (Fig. 6B-C). To assess more fully the changes in immune cell types after HSF1 knockdown, a control and HSF1 knockdown tumor was digested and subjected to single cell RNA sequencing (scRNA-seq). While shCTL 4T1 tumors were primarily comprised of tumor cells (71%) and macrophages (23%), the tumors expressing shHSF1 showed a more diverse population of cell types (Fig. 6D-F, Suppl. Fig. 8C-D). Specifically, HSF1 knockdown tumors showed increased proportions of several populations including CD8+ T cells (3.5 fold increase), B cells (14 fold increase), and neutrophils/granulocytes (13 fold increase) (Fig. 6F, Suppl. Fig. 8E). These data confirm a role for HSF1 in regulating changes of CD8+ T cells within tumors but also points towards a potentially larger immunologic switch in breast tumors catalyzed by a loss of HSF1 in cancer cells that appears to promote a robust immunogenic response.

**Figure 6:**
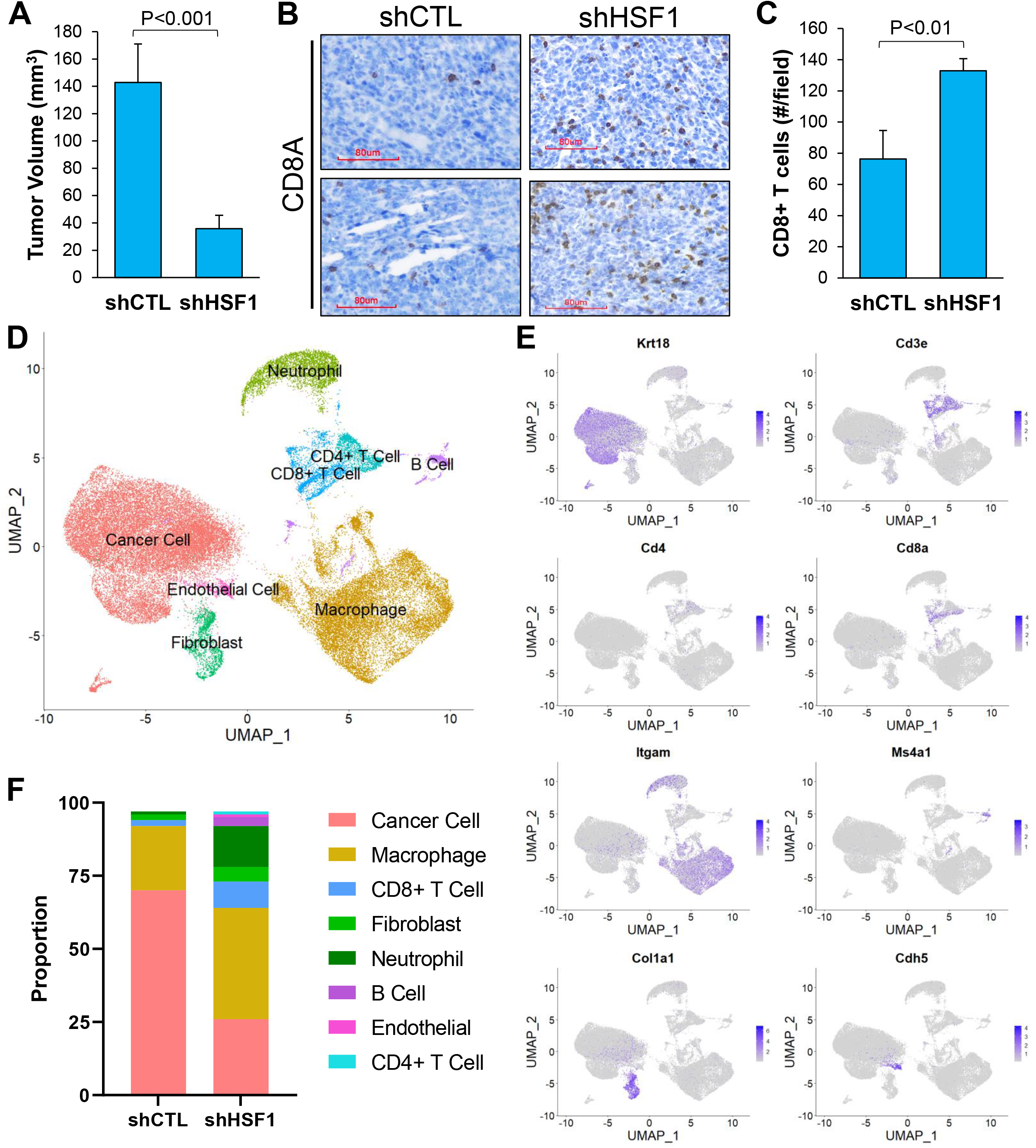
HSF1 Functionally Regulates the Amount of CD8+ T cells in Breast Tumors. A) 4T1 cells (5e4 cells) with (n=5) or without (n=5) HSF1 knockdown were grown orthotopically in Balb/c mice for 3 weeks. Tumor volume at the conclusion of the study is graphed. B) IHC was performed on tumors from (A) detecting Cd8a to identify CD8+ T cells. C) CD8+ T cells from (B) was quantified for control and HSF1 knockdown tumors by manual counting positive cells in >5 fields of the tumor tissue area. D-F) shCTL and shHSF1 tumors from (A) were subjected to scRNA-seq. Processed reads were used to map cell clusters for both samples using Seurat 4.2.0. The UMAP integrating both samples are shown in (D). These cell types were annotated using expression of specific marker genes for each population, for which a sample of these marker genes is shown in (E). The proportion of each cell population was also calculated and graphed in (F).

In addition to functionally regulating the attraction of CD8+ T cells to breast tumors, loss of HSF1 can intrinsically impact cancer cells. To determine the importance of CD8+ T cells to the loss of tumor volume after HSF1 knockdown, both control and HSF1 knockdown 4T1 cells were grown orthotopically in Balb/c mice with or without CD8+ T cell depletion. Splenic T cell counts confirmed that CD8+ T cells were indeed depleted at the end of the 3-week tumor growth period (Fig. 7A). Depletion of CD8+ T cells led to an expected increased volume of control tumors as CD8+ T cell depletion removes a tumor suppressor (Fig. 7B). Additionally, the depletion of CD8+ T cells rescued tumor growth with HSF1 knockdown, indicated by a greater increase in tumor growth after CD8+ T cell depletion (~50 fold increase) compared to the increase in control tumors (~9 fold increase). The decrease in tumor volume with HSF1 knockdown was accompanied by an increase in CD8+ T cells while mice with CD8+ T cell depletion were largely void of these cells within the tumors (Fig. 7C). These results suggest that CD8+ T cells were a significant contributor to the decreased tumor volume with HSF1 knockdown and that HSF1 can protect breast tumors from immune-mediated killing.

**Figure 7:**
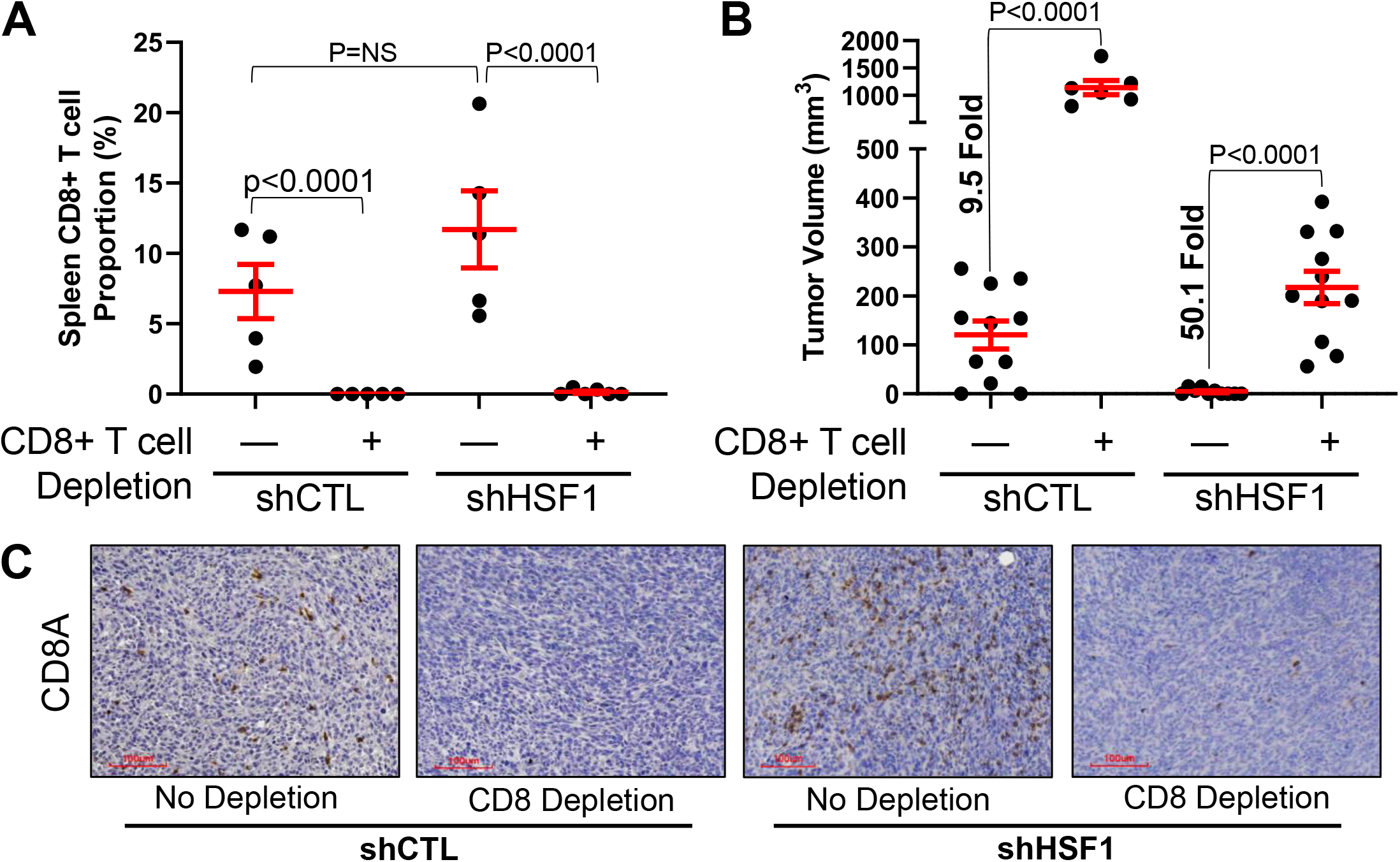
Depletion of CD8+ T cells Rescues Tumor Growth after HSF1 Knockdown. A) Balb/c mice were given either PBS control or CD8A antibodies to deplete CD8+ T cells in vivo. Control and HSF1 knockdown cells were then grown orthotopically for 3 weeks. Spleen were collected at the conclusion of the study, dissociated, and cells were subjected to flow cytometry to confirm the depletion of CD8+ T cells. B) Tumor volume at the conclusion of the study from (A) is plotted. C) Tumor tissue from (B) was subjected to IHC for CD8A to assess the CD8+ T cells.

### HSF1 suppresses expression and secretion of CCL5, which is necessary to attract CD8+ T cells after HSF1 knockdown

To understand how HSF1 activity in cancer cells results in suppression of CD8+ T cell infiltration into breast tumors, we tested whether HSF1 regulates expression or secretion of cytokines that attract CD8+ T cells. We compared expression of cytokines from RNA-seq of 4T1 control and HSF1 knockdown cells with a cytokine array detecting over 100 cytokines that was performed on conditioned media from 4T1 control and HSF1 knockdown cells. The only cytokine that showed a significant increase with HSF1 knockdown at the RNA level and secreted protein was CCL5/RANTES (Fig. 8A; Suppl. Fig. 9A), which is also a canonical chemoattractant cytokine for CD8+ T cells (39). The effect of HSF1 knockdown on CCL5 expression was confirmed with qPCR in both mouse and human breast cancer cells (Fig. 8B-C) while overexpression of HSF1 decreased CCL5 transcript levels (Suppl. Fig. 9B). HSF1 knockdown 4T1 tumors also showed an increase in CCL5 protein by IHC compared to control tumors (Fig. 8D). Additionally, assessing CCL5 levels by IHC in the same cohort of breast cancer specimens from Figure 5, CCL5 levels were significantly lower in tumors with high active HSF1, which remained significant when analyzing only TNBC patients (Fig. 8E-G).

**Figure 8:**
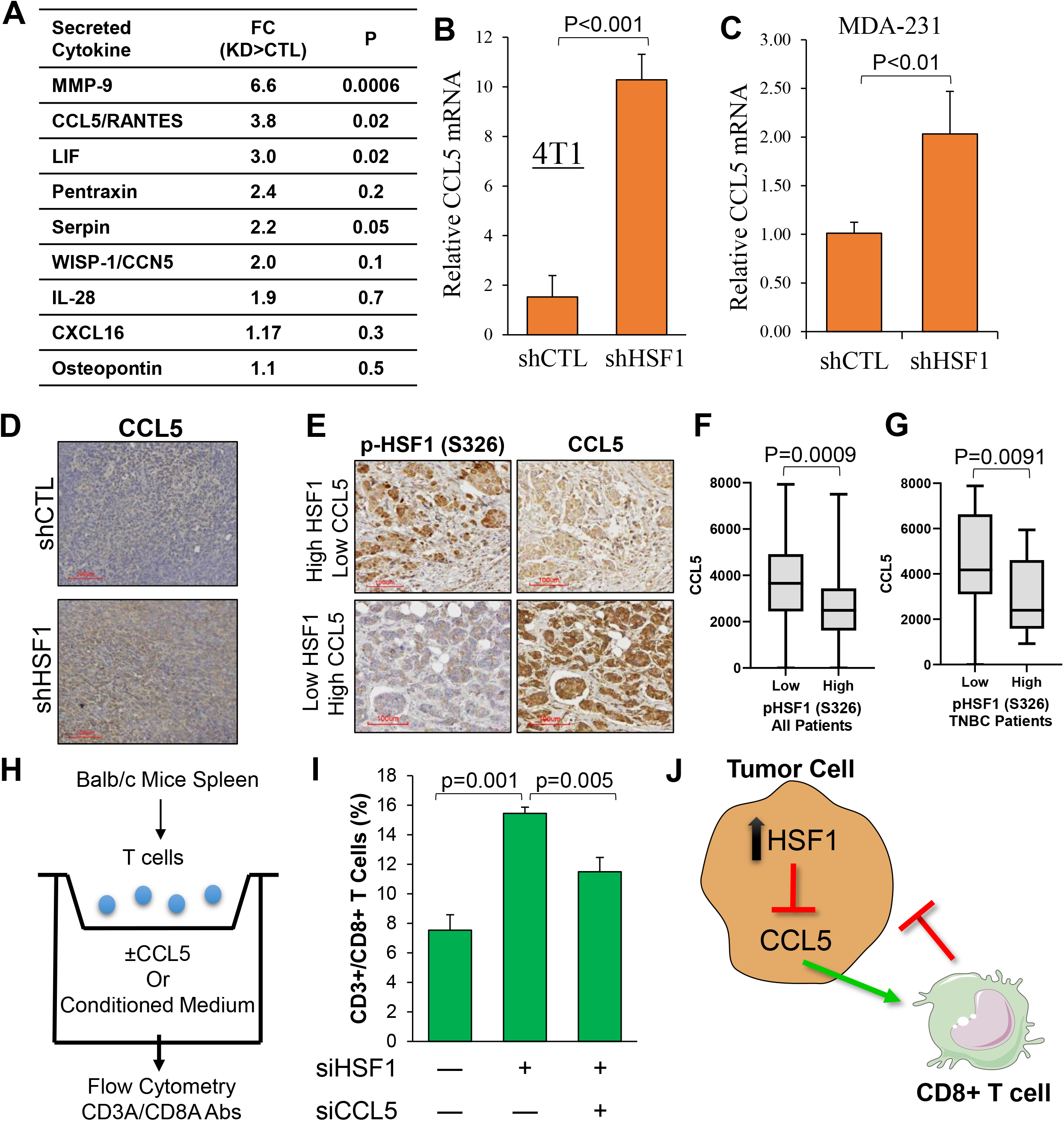
HSF1 suppresses CCL5 to prevent attraction of CD8+ T cells. A) Conditioned media was grown for 72 hours on control or HSF1 knockdown 4T1 cells. Conditioned media was then subjected to a cytokine array detecting over 100 cytokines. Cytokines are ordered by p value and fold change (FC) is calculated as HSF1 knockdown (KD) divided by control (CTL) cells. B-C) CCL5 mRNA levels in control and HSF1 knockdown 4T1 (B) and MDA-MB-231 (C) cells assessed by RT-qPCR. D) IHC of CCL5 in 4T1 control and HSF1 knockdown tumors from Fig. 6A. E) Tumor specimens from Fig. 5 were subjected to IHC for CCL5. CCL5 levels were quantified by QuPath. F) CCL5 levels are plotted in high (n=46) or low (n=68) active HSF1 tumors from all patients (n=114). G) CCL5 levels are plotted in high (n=21) or low (n=17) active HSF1 tumors from TNBC patients (n=38). H) Schematic of T cell transwell migration assay wherein pan-T cells were incubated in the upper chamber for 24 hours with conditioned media in the lower chamber, after which the lower chamber was subjected to flow cytometry to detect CD8+ T cells. I) Conditioned media from 4T1 cells expressing either control, HSF1, or HSF1+CCL5 siRNA were placed in the lower chamber for the T cell transwell migration assay. CD8+ T cell proportions are plotted for each group (n=5) and statistically compared using one-way ANOVA with Tukey’s posthoc test. J) Model indicating HSF1 suppresses CCL5 expression and secretion leading to decreased attraction of CD8+ T cells toward breast cancer cells.

To test the importance of increased expression and secretion of CCL5 after HSF1 knockdown, transwell migration of T cells was tested. In this assay, CD3+ T cells were isolated from Balb/c mice spleens and T cells were placed in the upper chamber while the lower chamber contained chemokines or conditioned media and incubated for 24 hrs followed by assessment of the lower chamber by flow cytometry for CD3+/CD8+ T cells (Fig. 8H). Adding exogenous CCL5 to the lower chamber resulted in a directed migration of CD8+ T cells to the lower chamber indicating the assay is responding to a positive control (Suppl. Fig. 9C). To test the effect of HSF1 knockdown, conditioned media from 4T1 cells with control or HSF1 siRNA were added the lower chamber of the T cell migration assay, which resulted in a significant increase in CD8+ T cell migration with HSF1 knockdown (Fig. 8I). Knockdown of both HSF1 and CCL5 significantly reduced the CD8+ T cell migration observed with only HSF1 knockdown (Fig. 8I), suggesting the increased expression and secretion of CCL5 after HSF1 knockdown is necessary to attract CD8+ T cells. Taken together, these data suggest a model whereby tumors with high activity of HSF1 leads to suppression of CCL5 transcript and secreted protein that ultimately prevents attraction of CD8+ T cells toward breast tumors allowing the tumors to evade immune-mediate destruction (Fig. 8J).

## Discussion

The major finding from these studies is activity of the stress response transcription factor HSF1 in breast cancer cells, known to support malignancy, prevents CD8+ T cells from attacking breast tumors. Furthermore, depletion of HSF1 resulted in a recruitment of CD8+ T cells that was found to be critical to reducing tumor volume after HSF1 depletion, thereby directly connecting HSF1 function to CD8+ T cells within breast tumors. The recruitment of CD8+ T cells appears to be part of an entire reprogramming of the tumor microenvironment after loss of HSF1 in cancer cells that was accompanied by increases in several immune populations such as neutrophils/granulocytes, B cells, and CD4+ T cells. While a direct connection between HSF1 function within tumors and immune cell presence or infiltration has not yet been established, a negative association between CD8+ T cells proportion estimates and HSF1 gene expression has been observed in breast cancer (32). The current study clearly indicates loss of HSF1 decreases tumor volume in an immune competent system, consistent with previous results (25) and point to HSF1 functioning as an immune-suppressive factor in the context of cancer. These data indicate for the first time that CD8+ T cells, specifically, are critical to the tumor suppressive effect of HSF1 depletion. We suggest that therapeutic targeting of HSF1 in breast tumors could lead to a reprogramming of the tumor microenvironment to be more immunogenic, which would enhance therapeutic response to several standard of care breast cancer therapies including taxanes and anthracyclines (11,40).

HSF1 has been known as the master regulator of the heat shock response since the mid-1980s. It was first observed to be altered in metastatic prostate cancer and has since been found to play a pleiotropic role in cancer regulating many functions in cancer cells from metabolism to proliferation. Due to the complex activation of HSF1 protein activity, the transcript levels have poor utility in predicting or assessing HSF1 activity. We identified a gene signature, named HAS, comprised of 23 genes that are direct HSF1 gene targets and depend on HSF1 for their expression. Increased HSF1 activity largely results in increased expression of the majority of the 23 genes, indicated by the high intra-gene correlation amongst these 23 genes providing for the first time an accurate and sensitive readout of HSF1 activity using transcript data. Studies wherein HSF1 gene was knocked down or exposed to heat shock, both of which have predictable effects on HSF1 activity, confirmed this observation. There is one previously reported HSF1 gene signature, the CaSig, developed from elegant studies identifying the many roles for HSF1 in cancer cells (19). The performance of the HAS in our studies is possibly due to the intra-gene correlation as the CaSig (456 genes) had low intra-gene correlation. Including intra-gene correlation as a criteria is an additional novel aspect to the HAS and the development of gene signatures. The application of the HAS will allow for future analysis of HSF1 transcriptional activity with many cancer-related studies that cannot be done based on HSF1 expression alone.

Our results indicate the influx of CD8+ T cells after HSF1 depletion is due to an increase in CCL5 expression and secretion, suggesting CCL5 would serve a tumor suppressive function. However, the role of CCL5 in breast cancer is controversial. A recent report indicates CCL5 is associated with breast cancer metastasis (41), while another study indicated therapeutic response to HDAC inhibitors in lung cancer is accompanied by a decrease in MYC function and an increase in CCL5 that supported therapeutic efficacy (42), suggesting tumor suppressive effects of CCL5. The disparity in results for the function of CCL5 in cancer is puzzling but is possibly related to expression of the CCL5 receptor, CCR5, on cancer cells. There is also precedent for an increase in cytokine-related signaling with HSF1 inhibition in other contexts, such as LPS exposure (43,44). Because these results saw a partial loss of CD8+ T cell attraction by silencing CCL5, it is likely that attraction of T cells after HSF1 depletion is regulated by additional mechanisms, including antigen presentation (45). Future studies will address other possible mechanisms by which HSF1 regulates the composition of the tumor microenvironment and the regulation of the CCL5 gene as it was not identified as a direct HSF1 target when generating the HAS.

Further investigation is needed to understand if immune suppression is a general feature of HSF1 biology. Inversely to cancer cells, neurodegenerative diseases share etiologies with an imbalance of proteostasis leading to protein aggregate formation in neurons. Opposite to cancer cells, these neurons lose HSF1 expression and activity with aging that ultimately escalates these diseases (46). Furthermore, several neurodegenerative diseases have been linked to immune-mediated mechanisms for neuronal death in recent years (47). Consequently, it is possible HSF1 serves to protect over-stressed cells from immune-mediated destruction and our findings point to how cancer cells have co-opted this function for their benefit and survival.

These studies indicate a novel mechanism regulating CD8+ T cell presence in breast tumors wherein hyperactive HSF1 suppresses CCL5 leading to decreased attraction of CD8+ T cells. Breast cancer is a low immunogenic tumor, which could partially be due to high basal activity of HSF1. Further studies will need to identify other mechanisms contributing to the effect of HSF1 on CD8+ T cells and other immune cells in the tumor microenvironment. Studies are underway to test inhibition of HSF1, with compounds such as DTHIB (48), can result in an influx of CD8+ T cells that would support targeting HSF1 as a therapeutic approach to making breast tumors more immune cell-rich. The role of HSF1 in CD8+ T cell activity or exhaustion warrants further investigation considering HSF1 has been reported to upregulate PD-L1 expression on cancer cells (49). These results generate several new lines of investigation for the role of HSF1 in new aspects of tumor biology and tumor-immune interactions.

## Materials and Methods

### Cell Culture

MDA-MB-231 (DMEM, Gibco) and 4T1 (RPMI, Gibco) cells were obtained from ATCC and were maintained at 37°C in 5% CO_2_ and culture media was supplemented with 10% fetal bovine serum (Corning) and 1% penicillin/streptomycin (Gibco). Cells were tested monthly for mycoplasma contamination using MycoAlert kit (Lonza). Lentiviral plasmids carrying control (5’-CCTAAGGTTAAGTCGCCCTCG-3’) or HSF1 (5’-GGAACAGCTTCCACGTGTTTG-3’) shRNA were synthesized by VectorBuilder using U6-driven promoter.

### RNA-Sequencing and Datasets Used

Total RNA from MDA-MB-231 and 4T1 cells were collected (ThermoFisher) and subjected to mRNA-sequencing using an Illumina HiSeq 4000. Datasets from the public domain were accessed from NCBI GEO (Suppl. Table 1). The TCGA-BRCA dataset was accessed through the TCGA data portal. The METABRIC dataset was accessed through the European Genome-Phenome Archive (EGA) under study ID EGAS00000000083.

### Integrated co-expression analysis

Integrated co-expression analysis was used to identify gene modules with strong correlation patterns across multiple datasets. Genes were examined that were direct targets of HSF1 and had decreased expression with HSF1 knockdown. We included 11 cancer expression datasets (Suppl. Table 2) in the integrated analysis. Raw .CEL files of the datasets were downloaded, processed, normalized using MAS5, and calculated the correlation matrix for the filtered genes. A binary matrix was constructed for each dataset where 1 indicated significant correlation and 0 indicated non-significant correlation. An average matrix for the 11 binary matrices was calculated, which was binarized such that only entries larger than a threshold *α* were assigned a value of 1. This binarized matrix was treated as an adjacent matrix for the genes and applied a greedy algorithm to identify the inter-connected modules. Here, we set *α* to be 0.3. Noted, we conducted robustness analysis for different values. The resulting gene module are highly overlapping. In total, we identified nine different modules.

### Heat Map Generation, Principal Component, and Gene Ontology (GO) Analysis

Z-scores were used to generate heat maps using Morpheus. Gene expression values for the respective genes within each signature were subjected to principal component analysis using GraphPad Prism 9. PC1 scores for samples from respective groups were used to rank order samples for survival analysis or used to compare signature scores between groups. Gene ontology for the HAS or genes associated with the HAS was performed using ShinyGO version 0.76. Genes associated with the HAS were selected through Pearson correlation of HAS score with all genes within the BRCA TCGA cohort.

### Survival Assessment

Patient survival was assessed using Kaplan-Meier plots within Prism 9. Log-Rank tests were used for determination of statistical differences between groups. Cox proportional hazard ratios were computed using PC1 scores from gene signatures with survival using SPSS 28.0 and calculated 95% confidence interval and computed p-values.

### Gene Set Enrichment Analysis (GSEA)

GSEA was done as previously described (20). Gene Cluster Text file (.gct) was generated from the TCGA BRCA, METABRIC, or GSE47561 breast cancer cohorts. Categorical class file (.cls) was generated by separating patients in these datasets based on high and low HAS score. The Gene Matrix file (.gmx) was generated using published gene signatures for immune cell types. The number of permutations was set to 1000 and the chip platform for TCGA gene lists was used. Heat maps were generated using Normalized enrichment scores (NES).

### Immune Profile Estimations

Immune proportion estimates were completed using deconvolution algorithms from TIMER, CIBERSORT, QuantiSeq, MCPCounter, and XCell for the TCGA-BRCA, METABRIC, and GSE47561 datasets using the TIMER2.0 portal (http://timer.cistrome.org). CD8+ T cell proportions were compared between high and low HAS groups using students t-test.

### Immunohistochemistry (IHC)

IHC was performed on a breast cancer tumor microarray (BR1141A; tissuearray.com). Antibodies used for IHC include pHSF1 (S326) (Abcam), CD3A (Santa Cruz), CD8A (CST), and CCL5 (Invitrogen). IHC was performed as previously described (20). Slides were imaged with Motic EasyScan scanner and analyzed with QuPath software (50).

### Animal Studies

All animal studies were performed under an approved institutional animal care and use committee protocol on the Indiana University-Bloomington campus. Control or HSF1 knockdown 4T1 cells (3e5) were injected into the mammary gland of 4-week old Balb/c mice and allowed to grow for 3-5 weeks. Tumor volume was measured with calipers. CD8+ T cell depletion was accomplished by administration of 200 ug of Cd8a (bioXcell) antibodies for 3 days and depletion was maintained throughout the study with 200 μg antibody injections (twice per week). CD8+ T cell depletion was confirmed at the end of the study by collecting the spleens and analyzing by flow cytometry.

### Single cell RNA-sequencing

Tumors from 4T1 shCTL or shHSF1 were excised then dissociated using a tumor dissociation kit (Miltenyi Biotech). Approximately 10,000 cells per sample with greater than 70% viability were used as input to the 10X Genomics Chromium system using the Chromium Next GEM Single Cell 3’ Kit v3.1. Libraries were sequenced using a NovaSeq 6000 with a NovaSeq S2 reagent kit v1.0 (100 cycles). Count matrices were generated with 10X Genomics Cell Ranger (v4.0.0) with default settings and genome assembly with GRCh38 was used. Resulting matrices were processed and analyzed in Seurat (v.4.2.0). Quality control and filtering removed low-quality cells and genes. The two samples were integrated using function ‘IntegrateData’ in Seurat. We then selected variable genes and performed uniform manifold approximation (UMAP) dimensionality reduction and cell clustering on the integrated data. Cluster markers were overlapped to canonical cell type-defining signature genes. Ultimately, we recovered 8 unique cell types from the two samples.

### Flow Cytometry

Single cell suspensions were subjected to incubation with mouse-specific Cd3a (Miltenyi) and Cd8a (Miltenyi) antibodies in series followed by incubation with Zombie viability dye (Invitrogen). After labelling, flow cytometry was performed using a MACS Quant (Miltenyi) system and data were analyzed with FlowJo 10.8.

### Cytokine Array

Control and HSF1 knockdown 4T1 cells were incubated in fresh media over 72 hours. Culture media was collected, centrifuged, and subjected to the Mouse XL Cytokine Array (R&D Systems) according to manufacturer’s instructions in triplicate.

### RT-qPCR

Total RNA from was extracted (ThermoFisher) and subjected to reverse transcription using RT Master Mix (Applied Biosystems). qPCR was performed with SYBR Green Master Mix (Applied Biosystems). qPCR primers used are listed in Supplemental Table 3.

### Transwell Migration of T cells

The assay was performed using a chemotaxis chamber (NeuroProbe) with 5 μm pore size filters. Briefly, the spleens were extracted from healthy Balb/c mice, dissociated using a gentleMACS and Spleen Isolation Kit (Miltenyi), and T cells isolated using a pan-T cell Isolation Kit (Miltenyi). T cells (1e6) were placed in the upper chamber in serum-free medium while the lower chamber contained serum-free medium with or without exogenous CCL5 or conditioned medium from 4T1 cells with pre-designed control siRNA, Hsf1 siRNA, or Ccl5 siRNA (Bioneer). T cells were incubated for 24 hours and subjected to flow cytometry with Cd3a/Cd8a antibodies with 5 biological replicates.

### Statistical Analysis

All statistical tests were performed as two-tailed tests. For two-group comparisons, students t-test was used. For multiple group comparisons, ANOVA with Tukey’s post-hoc test was used. All laboratory experiments were completed with a minimum of three biological replicates (e.g. qPCR, cytokine array) with flow cytometry experiments using a minimum of four biological replicates. Animal studies were completed with a minimum of five mice per group.

## Acknowledgements

This publication was made possible, in part, with support from the National Cancer Institute (K22CA207575, RLC; R01CA248033, XL), the Catherine Peachey Fund (RLC), and the Indiana Clinical and Translational Sciences Institute (RLC) funded, in part by Grant Number UL1TR002529 from the National Institutes of Health, National Center for Advancing Translational Sciences, Clinical and Translational Sciences Award. Research funding was also provided by P30CA82709-20 (Tumor Microenvironment and Metastasis Program) and Van Andel Institute through the Van Andel Institute – Stand Up To Cancer Epigenetics Dream Team (RLC, KPN). Stand Up To Cancer is a division of the Entertainment Industry Foundation, administered by AACR. The content is solely the responsibility of the authors and does not necessarily represent the official views of the National Institutes of Health. We would also like to acknowledge the Light Microscopy Imaging Center, the Flow Cytometry Facility, the Laboratory Animal Resources facility, and the Center for Medical Genomics at Indiana University for use of their core facilities.

## Author Contributions

RLC and CJ conceived the idea, oversaw completion of experiments, and analyzed the data. SC conceived of and executed many of the computational analyses. XL (along with RLC, SC) analyzed single cell RNA sequencing data along with conception of several included experiments. GS (with RLC) analyzed all histological tumor samples. KPN (along with RLC) helped with conception of breast cancer-related studies and analyses. SS, WCL, HR, and JW all completed multiple experiments and their associated analysis that are within the manuscript. RLC and CJ wrote the manuscript with contributions from SC, XL, and KPN.

