## Supplemental Figures for "HSF1 excludes CD8+ T cells from breast tumors via suppression of CCL5"

### SUPPLEMENTAL FIGURES AND LEGENDS

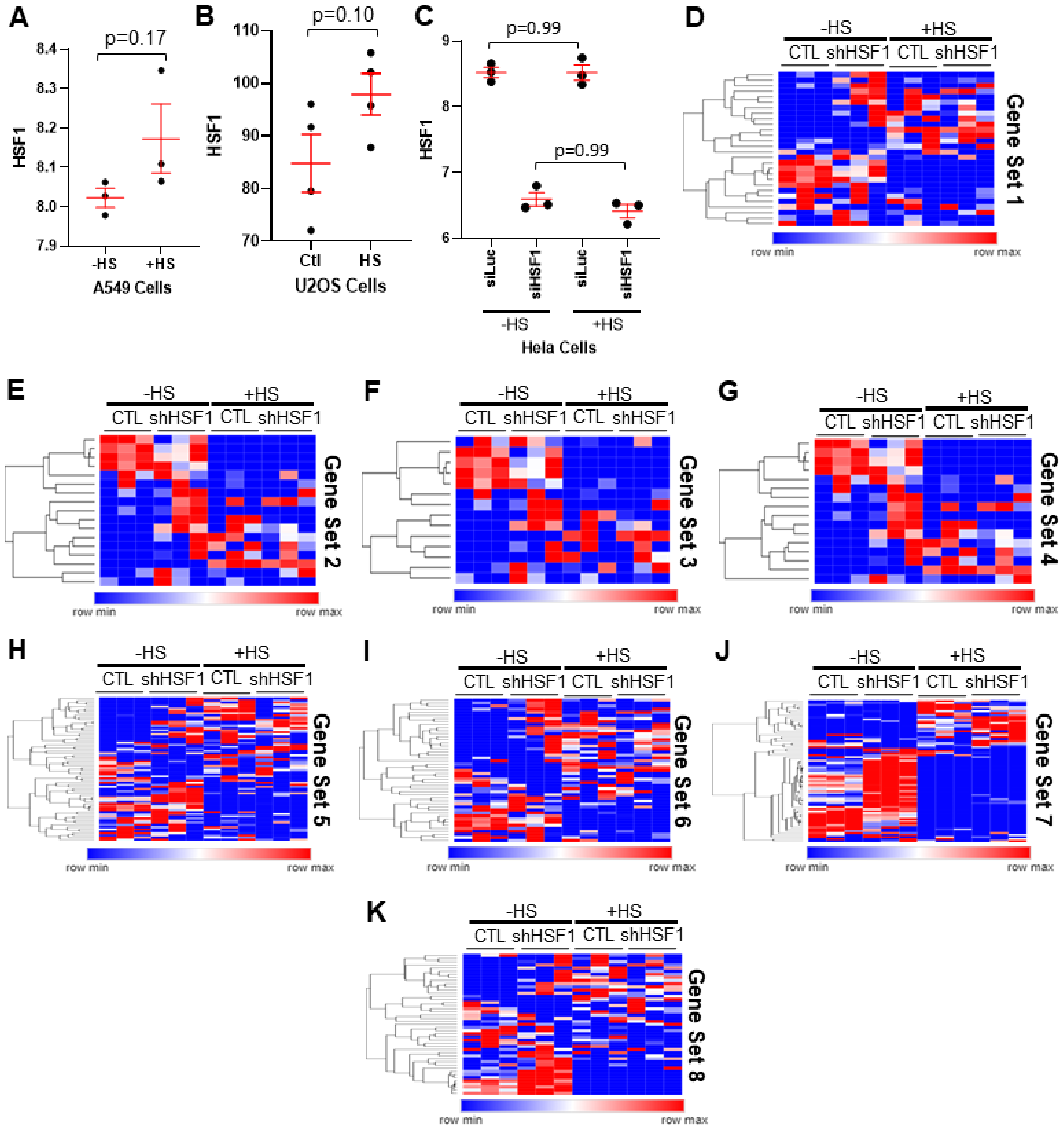


**Supplemental Figure 1: HSF1 activity gene sets in response to heat stress and HSF1 knockdown.** A-C) Indicated cell lines were subjected to heat shock or HSF1 knockdown and microarray performed to assess gene expression. Data from (A) was used from GSE83844. Data from (B) was used from GSE115973. Data from (C) was used from GSE3697. D-K) Each panel represents the expression of 8 unique gene sets identified as possible HSF1 activity signature gene sets. Data indicate how each gene set changes in response to HSF1 knockdown, heat stress, or both in Hela cells (GSE3697).

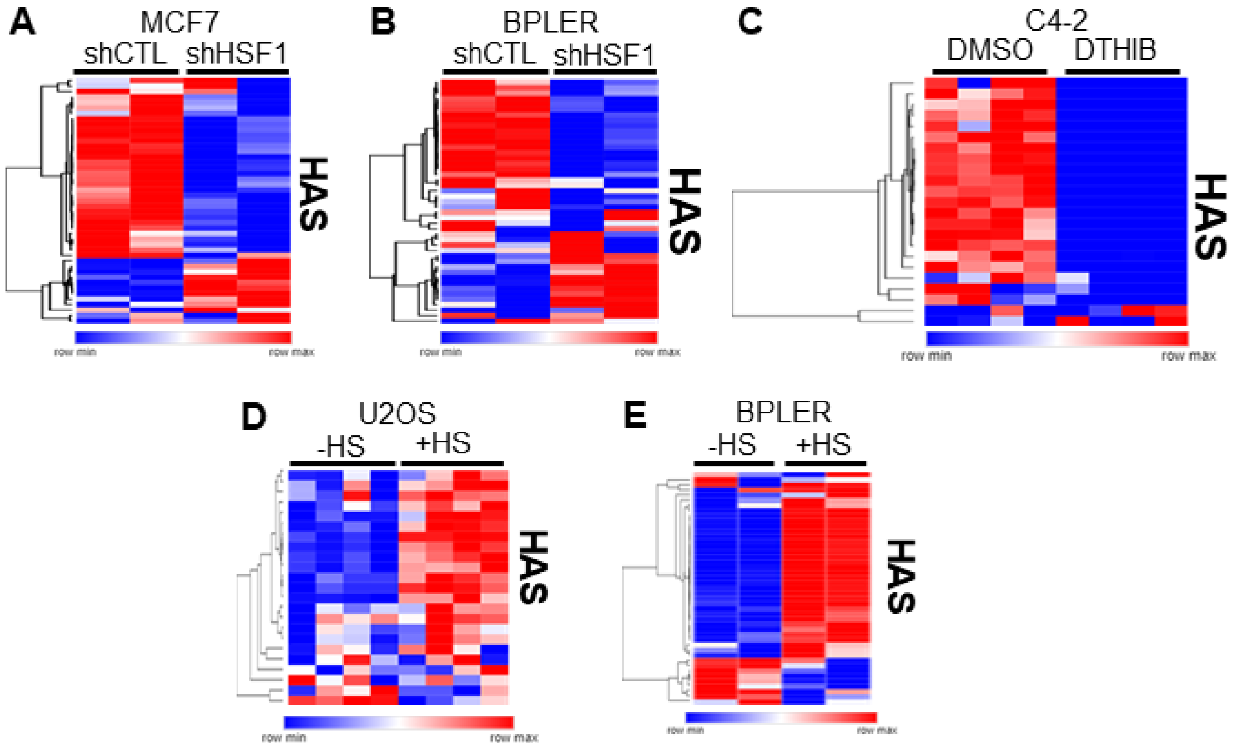


**Supplemental Figure 2: Response of HAS gene set to HSF1 knockdown or heat stress across cell lines.** A-B) Indicated cell lines were subjected to HSF1 knockdown and microarray to assess expression changes. Plots indicate the genes within the HAS and their response to HSF1 knockdown (GSE38232). C) C4-2 prostate cancer cells were treated with DTHIB or DMSO control and subjected to RNA-sequencing (GSE155248). Heat map indicates response of the HAS to HSF1 inhibition with DTHIB. D-E) Indicated cell lines were subjected to heat stress, followed by microarray to assess expression (GSE115973, GSE38232). Heat maps indicate response of the HAS to heat stress in these cell lines.

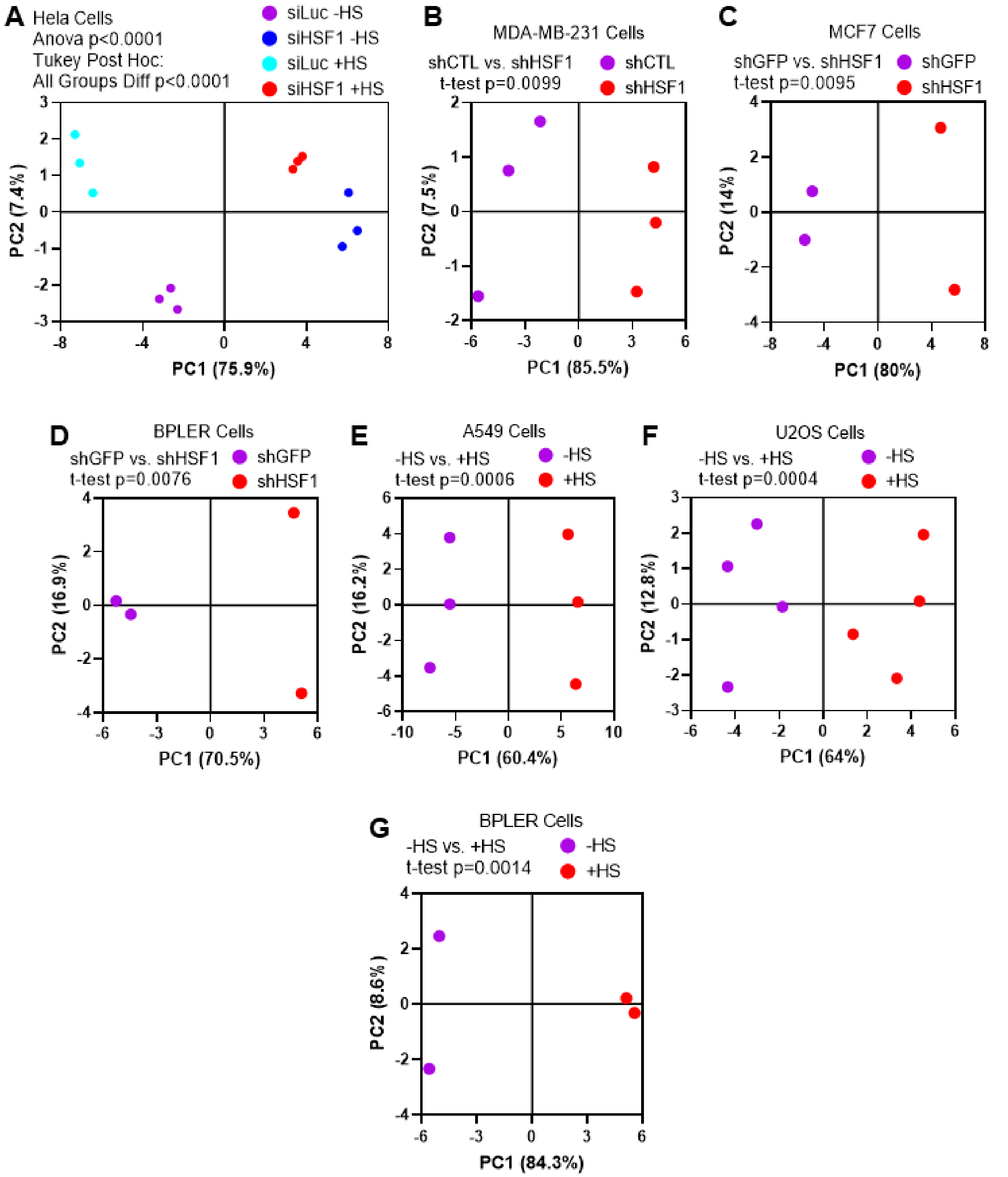


**Supplemental Figure 3: Principal component analysis of the HAS in response to HSF1 knockdown or heat stress.** A-G) Principal component analysis (PCA) for HAS genes was performed for all of the cell lines and samples indicated. PCA dimensional reduction maps are shown for each sample, indicating either knockdown of HSF1 or heat stress clearly separates the HAS gene set with these perturbations. A t-test or one-way ANOVA, where appropriate, was performed on PC1 scores for each sample to identify significant differences of the HAS gene set between groups.

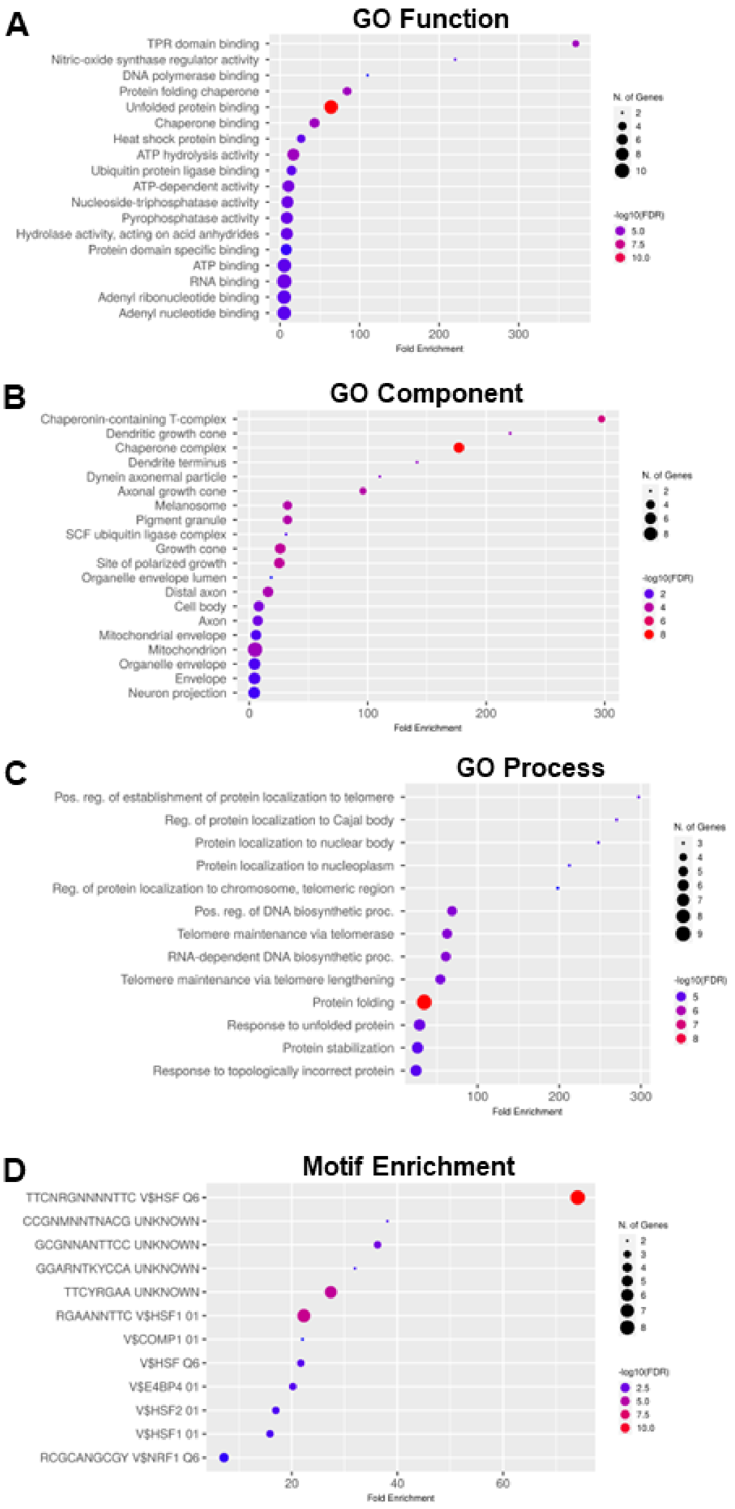


**Supplemental Figure 4: Gene ontology associated with the HAS gene set.** A-C) The TCGA BRCA cohort was used to identify the top genes correlated with the HAS. Gene ontology was performed using ShinyGO version 0.76. Lolliplot plots indicate ontologies most associated with the HAS. D) The 23 genes of the HAS gene set were analyzed for binding motif enrichment using ShinyGO version 0.76. The lollipop plot indicates motifs significantly enriched in the promoters of these genes.

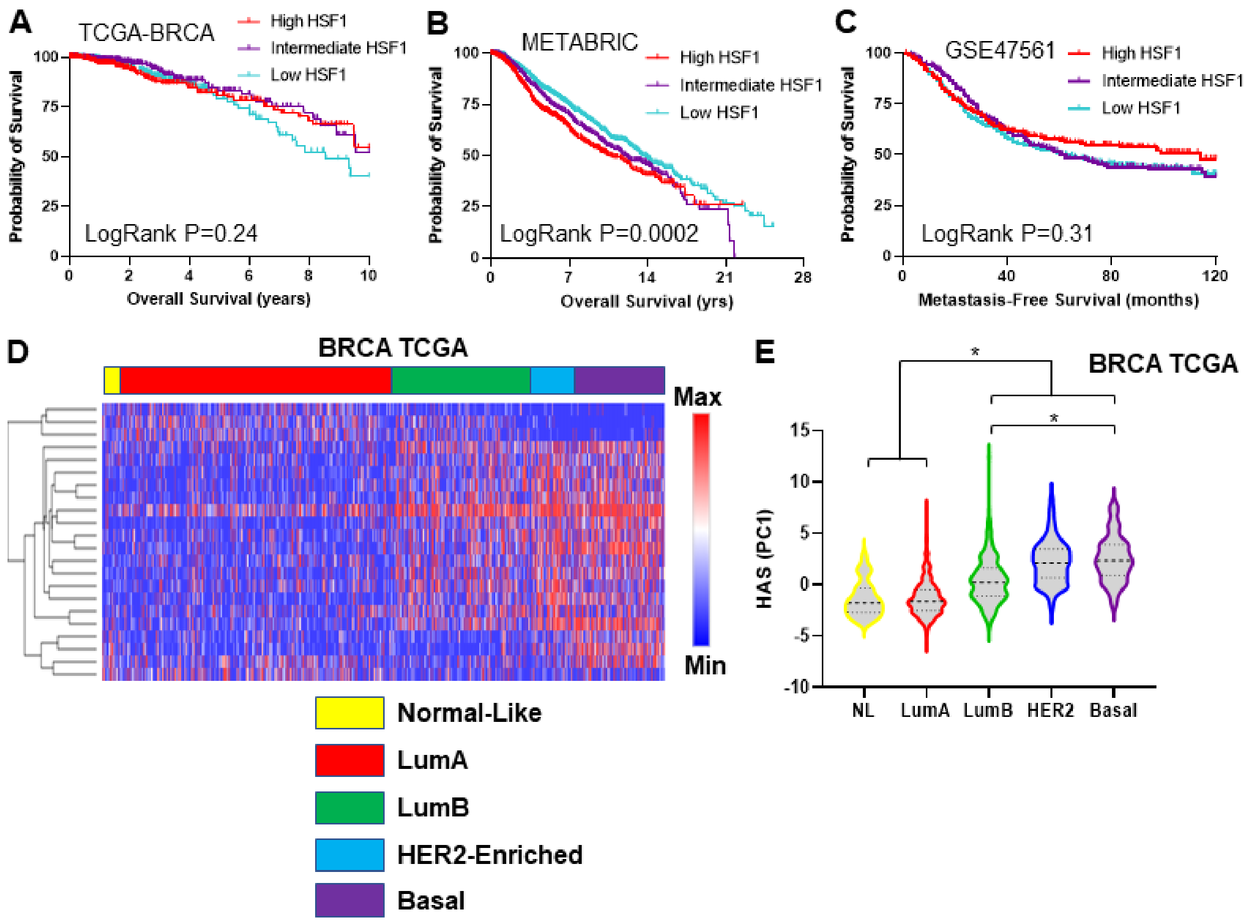


**Supplemental Figure 5: HAS and HSF1 association with breast cancer characteristics.** A-C) Expression of the HSF1 gene was used to separate patients into tertiles that were used to do Kaplan-Meier analysis of survival on the TCGA-BRCA cohort, the METBRIC cohort, and the breast cancer cohort GSE47561. P values are from Log Rank test of significance. D-E) The HAS gene set is plotted as a heat map for the TCGA-BRCA cohort separated by molecular subtypes (D). PCA of the HAS gene set in the TCGA-BRCA cohort was performed and PC1 scores plotted in (E). One-way ANOVA with Tukey's posthoc test was performed to test for significant differences between subtypes.

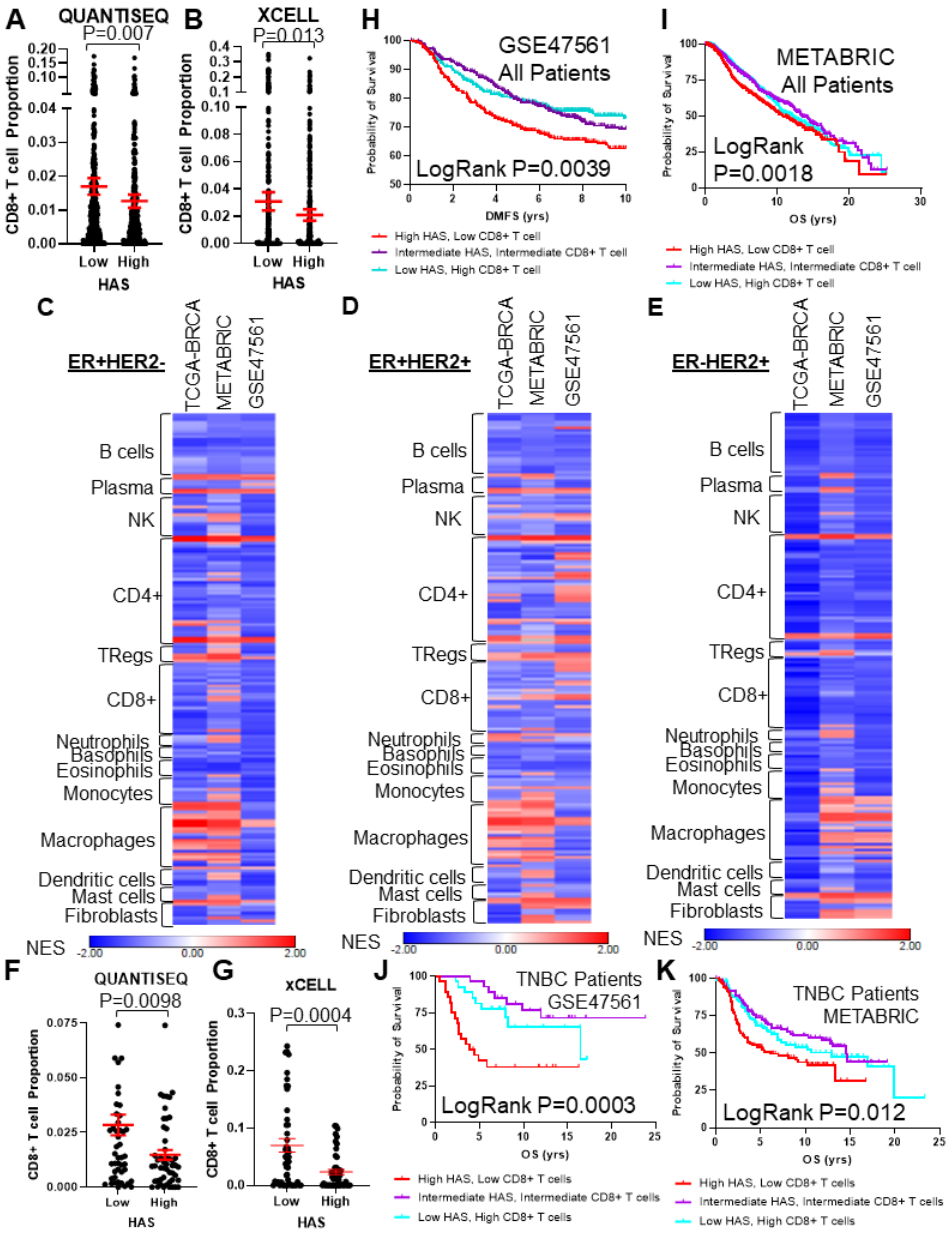


**Supplemental Figure 6: HAS association with immune cell types in breast cancer patient subpopulations.** A-B) CD8+ T cell proportions were estimated in the TCGA-BRCA cohort using the QuantiSeq (A) or XCELL (B) algorithm and compared between patients with high or low HAS. C-E) GSEA was performed in TCGA-BRCA, METABRIC, and GSE47561 cohorts for ER+/HER2- (C), ER+/HER2+ (D), and ER-/HER2+ (E) patients separated into high or low HAS scores. Signatures for immune cell types were assessed for enrichment with high or low HAS patients. Normalized enrichment scores (NES) are plotted on a heat map. F-G) CD8+ T cell proportions estimated from QuantiSeq (F) and XCELL (G) for only TNBC patients and were compared between high and low HAS tumors. H-I) All patients in the GSE47561 (H) or METABRIC (I) cohorts were separated by HAS scores and CD8+ T cell proportions estimated by CIBERSORT and Kaplan-Meier graphs were plotted for patient outcomes using all patients. J-K) Only TNBC patients in the GSE47561 (H) or METABRIC (I) cohorts were separated by HAS scores and CD8+ T cell proportions estimated by CIBERSORT and Kaplan-Meier graphs were plotted for patient outcomes using all patients.

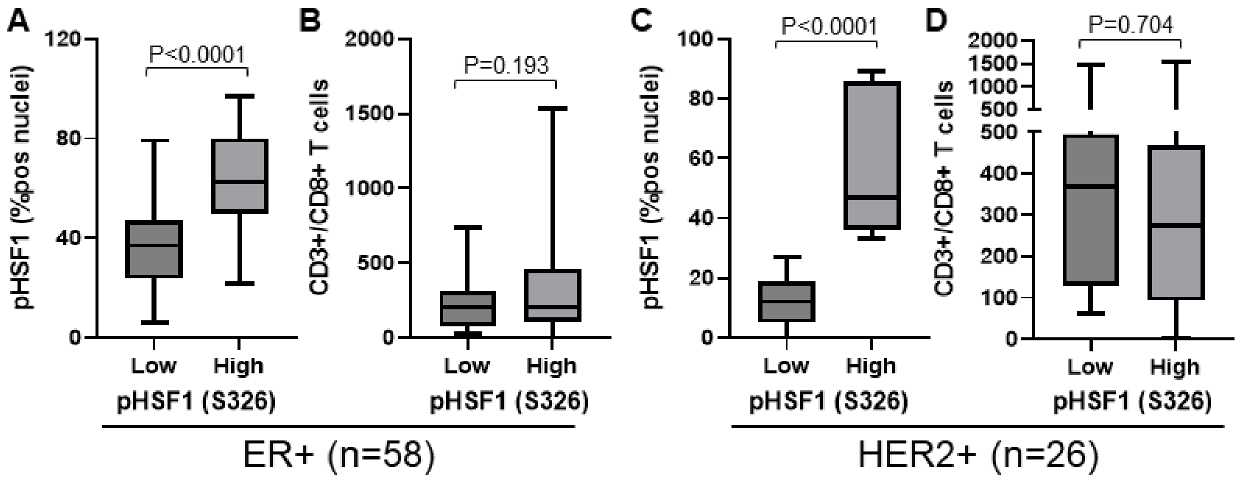


**Supplemental Figure 7: HSF1 activation and CD8+ T cells in human tumor specimens from ER+ and HER2+ tumors.** A cohort of 114 breast tumors were subjected to IHC for active HSF1 (pS326) and Cd3a/Cd8a antibodies to detect CD8+ T cells. A-B) Sub-analysis of only ER+ patients wherein tumors were separated into high (n=30) and low (n=28) HSF1 activity (A) and CD8+ T cells were compared amongst these groups (B). C-D) Sub-analysis of only HER2+ patients wherein tumors were separated into high (n=11) and low (n=15) HSF1 activity (C) and CD8+ T cells were compared amongst these groups (D). Students t-test was used to detect statistical differences.

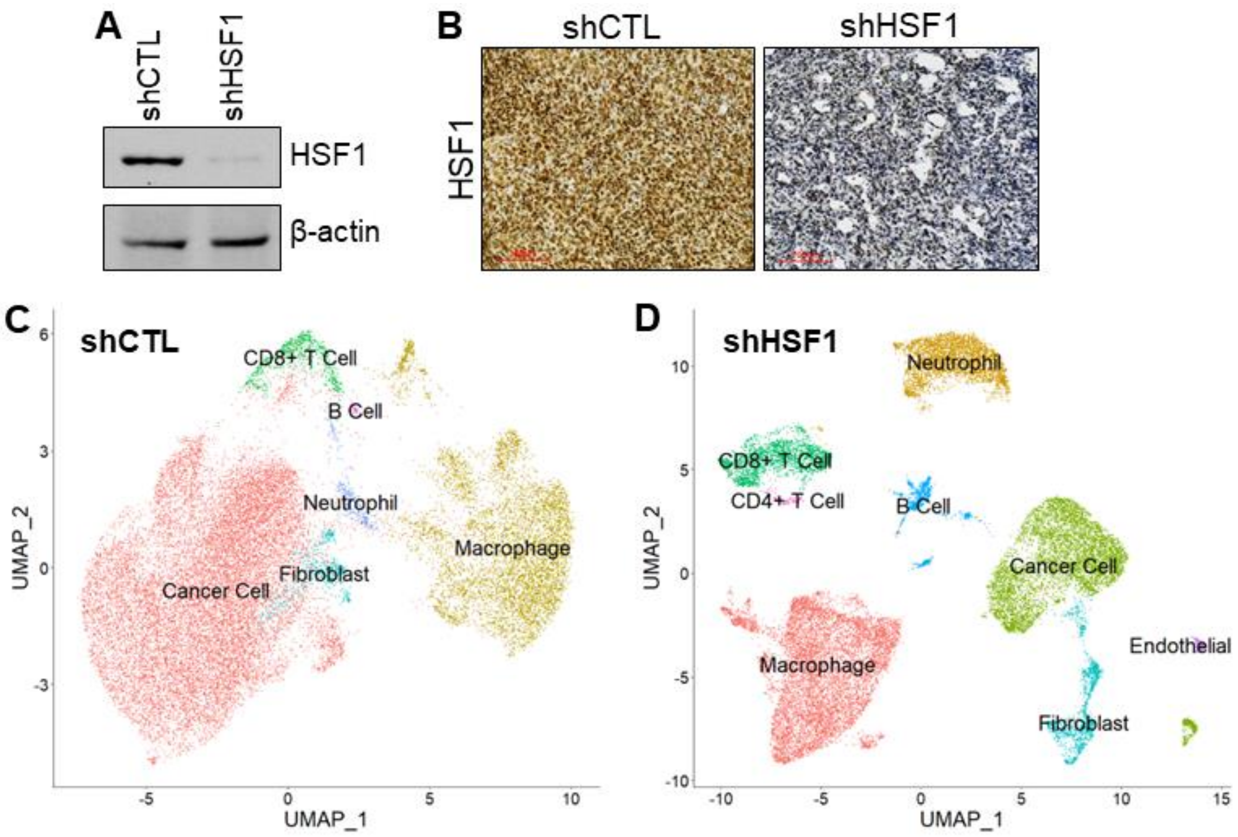


E

| Cell Type | shCTL | shHSF1 | Fold Change |
|---|---|---|---|
| Cancer Cell | 70.8 | 26.1 | 0.37 |
| Macrophage | 22.6 | 38.8 | 1.72 |
| CD8+ T cell | 2.84 | 9.77 | 3.44 |
| Fibroblast | 2.29 | 5.34 | 2.33 |
| Neutrophil | 1.13 | 14.8 | 13.1 |
| B cell | 0.27 | 3.73 | 13.9 |
| Endothelial | 0 | 0.008 | N/A |
| CD4+ T cell | 0 | 0.006 | N/A |

**Supplemental Figure 8: Knockdown of HSF1 restructures the tumor microenvironment.** A) 4T1 shCTL and shHSF1 stable cell lines were subjected to immunoblotting to assess efficacy of HSF1 knockdown. B) shCTL and shHSF1 cells were grown in Balb/c mice and tumors were extracted and subjected to paraffin-embedding. Unstained slides were subjected to IHC with HSF1 antibodies. C-D) shCTL and shHSF1 tumors from (B) were subjected to scRNA-seq. Processed reads were used to map cell clusters for both samples using Seurat 4.2.0. UMAPs for the shCTL tumor (C) and the shHSF1 tumor (D) are presented separately. E) Proportions of individual cell types based on analysis in (C-D) are listed, including the fold change of each cell type.

A

| Cytokine | FC (KD>CTL) | P |
|---|---|---|
| IL1a | 13.3 | 0.001 |
| CXCL16 | 3.38 | 0.010 |
| CCL5 | 4.12 | 0.019 |
| IL17f | 1.48 | 0.049 |
| CCL25 | 0.72 | 0.009 |

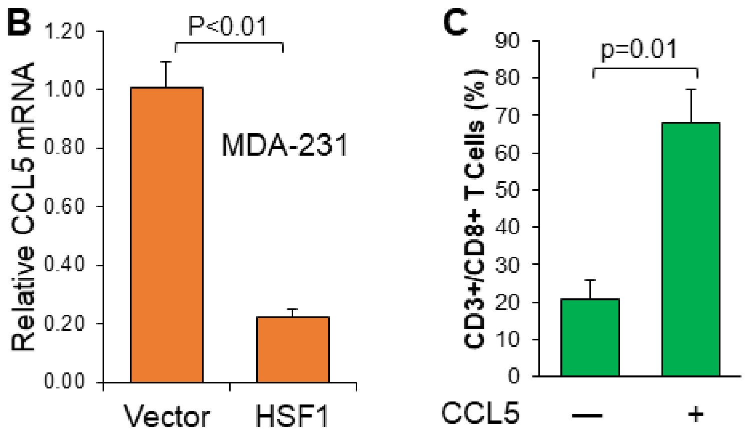


**Supplemental Figure 9: Identification of CCL5 as a downstream effector after knockdown of HSF1.** A) 4T1 shCTL and shHSF1 cells were subjected to RNA-seq. Cytokines were assessed for significant differences between these groups and those that were significant are listed, including their fold change and significance. B) Human MDA-MB-231 cells were transfected with empty vector or HSF1 and total RNA was subjected to RT-qPCR with CCL5 primers. C) Balb/c pan-T cells were incubated in the upper chamber of a transwell chamber for 24 hours in the presence or absence of exogenous CCL5. After 24 hours, the lower chamber was subjected to flow cytometry with CD3a/Cd8a antibodies. The proportion of CD8+ T cells between groups is plotted and significance was tested with a student's t-test.
